# Impact of various preservation and storage methods on the viability of mycoplasma field strains isolated in Mali

**DOI:** 10.1101/2021.10.21.465280

**Authors:** Amadou Sery, Cheick Abou Kounta Sidibe, Mamadou Kone, Bekaye Sacko, Joseph Awuni, William Amafu, Mamadou Niang

**Author notes:** Corresponding author (AS).

## Abstract

The survival of five mycoplasma strains was studied in different storage media (Mycoplasma complet media without cryopreservative agent, Mycoplasma complete media with addition of horse serum, Mycoplasma complete media with addition of glycerol and lyophilized cultures without stabilizer) under different temperatures (+37°C, +4°C, −20°C, −85°C) during 24 months. Five Mycoplasma strains, *Mycoplasma mycoides subsp mycoides* (Mmm), *Mycoplasma bovis* (Mb), *Mycoplasma agalactiae* (Ma), *Mycoplasma gallisepticum* (Mg) and *Mycoplasma synoviae* (Ms) were isolated from various parts of the country. The initial titers of the strains determined by the agar plate count before storage were 42.4×10^7^ UFC/ml (8.6 log UFC/ml) for Mmm strain; 32.4×10^8^ UFC/ml (9.51 log UFC/ml) for M.bovis strain; 12.4×10^9^ UFC/ml (10.09 log UFC/ml) for Ma strain; 2.4×10^9^ UFC/ml (9.38 log UFC/ml) for Mg and 2.8×10^9^ UFC/ml (9.45 log UFC/ml) for Ms strain. After 3 weeks of storage, no viable mycoplasmas were detected in all the conservation media at +37°C and after 3 months of storage at +4°C except for the lyophilized cultures in which an average viability rate of 17.81% was observed. Overall, the mycoplasma strains remained viable at freezing temperatures after 24 months regardless of the storage medium, but with decreasing titers, which was noticeable with mycoplasma complete media, and mycoplasma media with horse serum. Conversely, at −20°C the average viability rates after 24 months of storage were 84.36% (with glycerol) and 90.04% (lyophilized cultures). At −85°C after 24 months of storage, this was 87.98% (with glycerol) and 91.44% (lyophilized cultures). These findings suggest that, in the absence of the lysophylisation process, the addition of glycerol may be recommended for long-term storage of frozen mycoplasma isolates.

## Introduction

Mycoplasmas are bacteria that lack a cell wall, have demanding nutritional requirements and are characterized by causing diseases in plants, animals and humans and frequently contaminating cell cultures. The absence of a rigid cell wall makes them potentially vulnerable to environmental changes but they can nevertheless be preserved by low temperature storage, especially by freezing. Numerous studies have been carried out on the epidemiology and control of diseases caused by mycoplasmas (mycoplasmoses) by isolating the wild strains responsible for pneumonia, mastitis and arthritis [1–6]. However, very few studies have been conducted on methods of storing specimens and isolates of mycoplasmas using storage media and temperatures. The duration of the viability of mycoplasma strains under storage conditions depends on several factors, including the nature of the culture medium, as the protein content is probably responsible for the stability of the mycoplasmas over a long period [7] and also on temperature as well as addition of cryopreservatives agents such as glycerol and dimethyl sulfoxide (DMSO). During the storage of milk samples, the shelf life of *Mycoplasma agalactiae* and *Mycoplasma mycoides subsp. capri* was dependent on the temperature of refrigeration and freezing [8]. Cryopreservative agents are thought to improve the survival of *Mycoplasma bovis* in milk for both transport and long-term storage [9]. The critical effect of the process of thawing was evaluated with milk samples for the detection of mycoplasmas where thawing at room temperature was better than in a water bath at +37°C [10]. Similarly, with respect to the effect of the environment, strains of *Mycoplasma gallisepticum* and *Mycoplasma synoviae* remained viable for up to 2 to 4 days and 2 to 3 days, respectively [11]. Thus, appropriate storage conditions are necessary to achieve maximum survival of the mycoplasma strains.

This study evaluated the viability of *Mycoplasma mycoides subsp mycoides* (Mmm), *Mycoplasma bovis* (Mb) *Mycoplasma agalactiae* (Ma), *Mycoplasma gallisepticum* (Mg), *Mycoplasma synoviae* (Ms) using various storage media (Hayflick complete medium without cryopreservative agent, Hayflick with addition of horse serum and Hayflick with addition of glycerol) and lyophilized cultures of each strain after 24 months of storage under four different storage temperature ranges (+37°C, +4°C, −20°C and −85°C).

## Material and method

### Growth media and isolation of mycoplasma strains

Five strains of mycoplasma were obtained by isolation from lung tissue samples, pleural fluid, milk and inner ear swabs from cattle and small ruminants. Lung tissue samples were obtained in abattoirs after post mortem inspection. Routinely, the bleeding of animals in all abattoirs is carried out after mechanical stunning with a perforating rod gun (or matador). These were strains of Mmm, the causal agent for contagious bovine pleuropneumonia (CBPP), Mb, the agent responsible for pneumopathies and mastitis in cattle, Ma, the causal agent for contagious agalactia in young ruminants, Mg and Ms, responsible for avian mycoplasmosis.

These organisms were cultured in three specific media, which were Gourley′s medium (Bacto tryptose, Glucose, Sodium chloride, Sodium Phosphate Dissodique, Glycerol, Yeast Extract) [12] for the isolation of Mmm, the modified Hayflick medium [13] (Glucose, Sodium pyruvate) for Mbovis and Ma and Frey medium (Glucose, NAD, Thallium acetate) [14] for Mg and Ms. The base medium (PPLO broth, PPLO agar) was prepared in the proportion of 42 g of PPLO and 10 g of noble agar / liter then sterilized by autoclaving at 121°C for 15 min. As for the mycoplasma growth supplements composed of horse serum, fresh yeast extract, thallium acetate and ampicillin, they were sterilized by filtration through 0.45 μm then 0.22 μm. The complete mycoplasma medium was composed of basal medium (70%) and mycoplasma growth supplements (30%) (i.e. 70 ml of PPLO and 30 ml of growth supplement).

### *Mycoplasma mycoides mycoides* (Mmm) growth medium

Gourley′s medium (Gourlay et al., 1979) for *Mycoplasma mycoides mycoides* isolation: Bacto tryptose (BD / Ref 211713 / Batch 5299713), Glucose (SIGMA / Ref G5767 Batch BCBC0777), Chloride sodium (Aldrich / Ref 7647-14-5 / Batch 03609JT), Dissodium Sodium Phosphate (SIGMA / Ref 59763 / Batch BCBC3154), Glycerol (SIGMA / Ref 49770 / Batch BCBB7435), Yeast Extract (Fluka / Ref 70161 / Batch 0001439171), Horse serum (Gibco / Batch 1750660), Penicillin G (SIGMA / Ref 46609 / Batch BCBW2380); Noble agar (BBL / Ref 211456 / Batch 1112313).

### *Mycoplasma bovis* (Mbovis) and *Mycoplasma agalactiae* (Ma) medium

Modified Hyflick medium (Hayflick & Stanbridge, 1967) for culturing *Mycoplasma bovis* and *Mycoplasma agalactiae*: Glucose (SIGMA / Batch BCBC0777), Sodium pyruvate (Sigma / Batch 115K07251), Horse serum (Gibco / Batch 1750660), Fresh yeast extract (LCV-ELF / Batch 19005), Ampicillin (Sigma / Batch BCBR6229V), Thallium acetate (Alfa Aesar / Batch: G06Y019 / Ref: 11842).

### *Mycoplasma gallisepticum* (Mg) and *Mycoplasma synoviae* (Ms) growth medium

Dextrose (LAB M / Batch: Q23079 / Ref: 100743), Arginine (IGN / Batch: 3253E / Ref: 100743), Fetal calf serum (Eurobio / Ref: CVFSVF00-04 / Batch 531808), Horse serum (Gibco / Batch: 1750660 / Ref: 26050-088), Fresh yeast extract (Ref: ELF_LCV19005 / Batch: 001), NAD (Sigma / Ref: N3014 / Batch: 30K7068), Thalium acetate (Alfa Aesar / Batch: G06Y019 / Ref: 11842), Penicillin (Sigma / Ref: 46609 / Batch: BCBW2380).

### Preparation of mycoplasma suspensions for storage

Five (5) strains of pathogenic mycoplasmas were selected for their storage time in different storage media and temperatures. These local mycoplasma strains were: *Mycoplasma mycoides mycoides* (Mmm) (agent of contagious bovine pleuropneumonia (CBPP); *Mycoplasma bovis* (Mbovis) (agent of pneumonia and mastitis in cattle); *Mycoplasma agalactiae* (Ma) (agent of contagious agalactia in small ruminants); *Mycoplasma gallisepticum* (Mg) (responsible of avian mycoplasmosis); *Mycoplasma synoviae* (Ms) (*responsible for avian mycoplasmosis)*.

For cultivation, dilutions to 1/10th in eight (8) tubes (10^−1^–10^−8^) to 1 ml of sample in 9 ml of growth medium (broth) were carried out then 8 dishs of agar per sample were prepared (i.e. one agar dish per dilution from −1 to −8).

On each agar dish 100 μl of the dilution are deposited and spread on the surface of agar (100 μl of the dilution −1 on the agar plate labeled −1). After incubation at +37°C in humid atmosphere at 5% CO2, the dishes are examined every day for bacterial growth signs (change in color of the broth and the characteristic morphology of mycoplasma colonies “in egg on flat “).

The base medium (PPLO broth, PPLO agar) was prepared in the proportion of 42g of PPLO and 10g of noble agar/liter then sterilized by autoclaving at 121°C for 15 min. All the growth supplements composed of horse serum, fresh yeast extract, thallium acetate and ampicillin were sterilized by filtration through 0.45μm then 0.22μm. The complete mycoplasma medium was composed of 70% basal medium and 30% growth supplement (i.e. 70ml of PPLO and 30 ml of growth supplement).

Each diluted mycoplasma strain was aliquoted into 15 Eppendorf tubes (1ml each) on the same day of harvest. For total of 1300 tubes of 1ml stored, 60 tubes were at +37°C, 120 at +4°C, 560 at −20°C and 560 tubes at −85°C, i.e. 260 tubes of 1ml of each strain. At each titration, an aliquot of each storage medium was grown on selective media. At the same time, each mycoplasma strain was lyophilized in a volume of 2.5ml / vial produced in sufficient number for each storage temperature.

### Preparation of storage media

The following storage media were selected for this study: (a) Control medium without preservative additive (M1), (b) Complete medium for mycoplasmas as a preservative additive (M2), (c) Horse serum as a preservative additive (M3), (d) 60% glycerol as a preservative additive (M4) and (e) Lyophilisate culture without stabilizer as preservative (M5). For each preparation, 500µl of each strain were added to 500µl of each storage medium to have 1ml of final volume to store (50% and 50%, v/v).

### Determination of number of viable mycoplasmas

The effect of storage methods on the viability of mycoplasma strains was assessed by series of the agar plate count before (Day 0) and during (Week 1, Week 2, Week 3, Month 1 and Month 24) storage. The number of viable mycoplasmas was determined by the colony count method with several dilutions to 1/10 of the mycoplasma suspensions in proportion to 0.1 ml / agar plate. The dilutions giving 30 to 300 colonies were counted by means of a stereomicroscope and the results were expressed in colony forming units (CFU) per milliliter of suspension as follows:

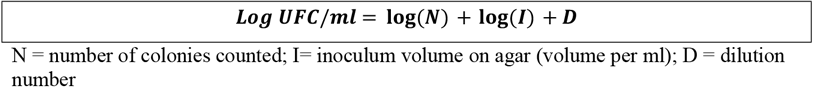

### Statistical analysis

Data was recorded and classified on the Excel spreadsheet software (Microsoft excel 10). The statistical tests used were the log means comparison (log CFU / ml) by strain, taking into account the fixed effect of media and storage temperatures (+37, +4, −20 and −85°C) by analysis of variance (ANOVA) with Stata 12.0 software and the proportions comparison (viability rate) by the chi2 test with R software (R i386 4.0.3). A significance level of 0.05 was retained for the comparison of various parameters.

### Ethical considerations

This study obtained the approval of the ethics committee of the « Institut National de Recherche en Sante’ Publique (INRSP) du Mali » following the decision N° 05/2019 / CE - INRSP. In addition, permission to carry out the study was obtained from the livestock owners. An informed consent document detailing the research objective and procedure was made available for them and a written informed consent was obtained. The Blood sampling procedures of animals were done with respect for animal welfare. In all abattoirs, mechanical stunning is a routine process for bleeding animals.

## Results

Overall, the mycoplasma strains remained viable at freezing temperatures after 24 months regardless of the storage medium. However, the number of viable mycoplasmas decreased slightly over time. The decrease was much smaller at storage temperature of −85°C than at − 20°C and in most cases, viability was higher in lyophilized cultures and glycerol than in horse serum and medium only (*p = 0*.*03*). On the other hand, the reduction in the viability of mycoplasma strains was much greater when they are stored at +37°C where the loss was maximum after 3 weeks (97.9%) and at +4°C with a maximum loss after 3 months of storage except lyophilized cultures (99.3%).

### Viability of mycoplasma strains at +37°C

The results obtained from this study showed that at +37°C, all the strains of mycoplasmas outside the incubation period were not viable beyond 3 weeks (Fig 1).

**Fig 1.**
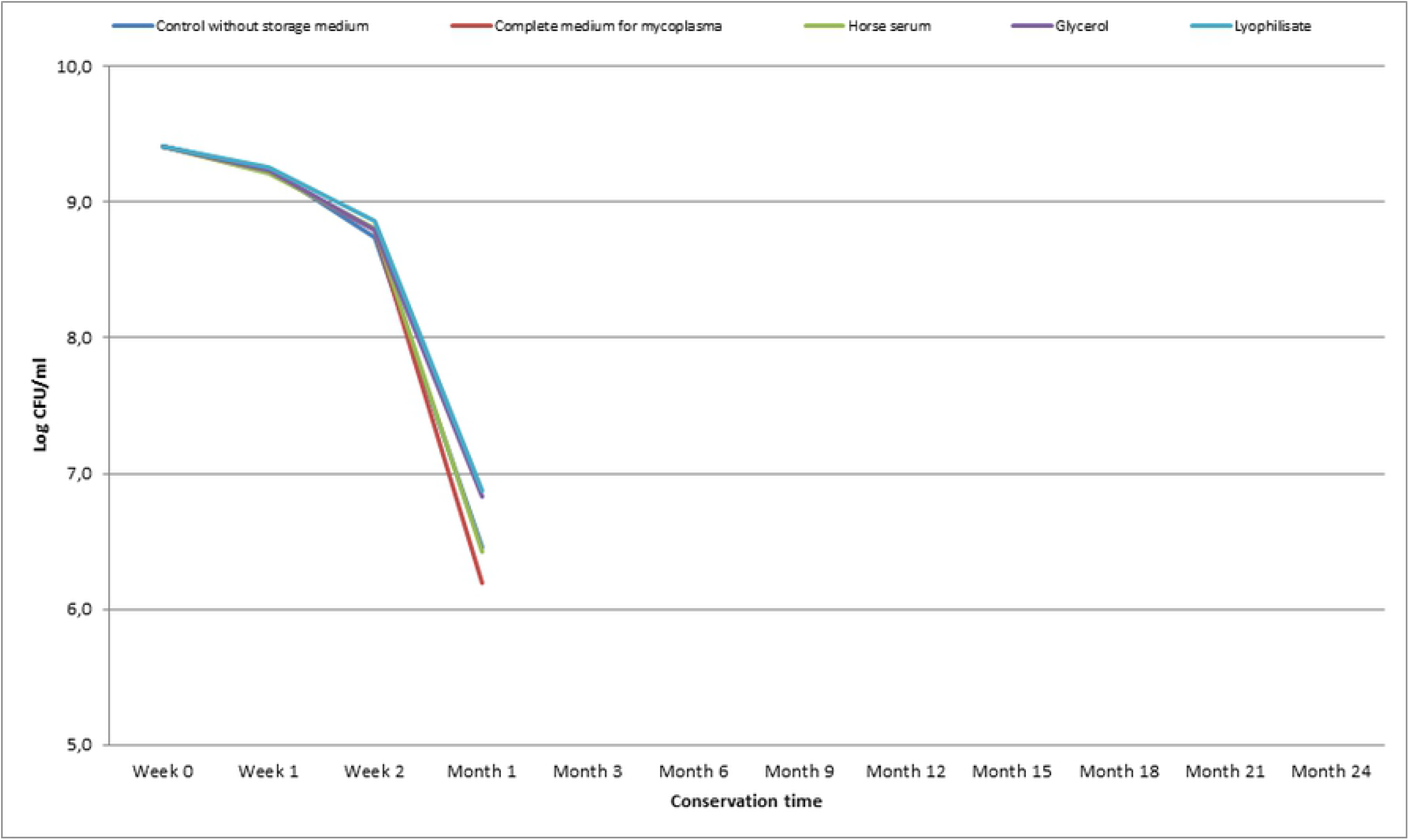
Mycoplasma viability at +37°C. Mean loss (in log) of the titre of 5 strains of mycoplasmas (Mmm, Mb, Ma, Mg, Ms) during 24 months of storage in different preservatives.

Overall, a very low non-significant overall mean viability (*p = 0*.*89*) was visible after 3 weeks of storage at +37°C with viability rates of 2.14%, (without preservative) at 4.38% (lyophilized cultures). A very low viability mycoplasma cultures was observed after 3 weeks of storage at +37°C.

The viability rates varied from 5. 91% (without preservative) to 12.98% (lyophilized cultures) for Mmm (*p = 0*.*42*), from 0.42% (without preservative) at 1.16% (lyophilized cultures) for Mb (*p = 0*.*95*), from 2.48% to 5.65%, respectively without preservative and the lyophilized cultures for Ma (*p = 0*.*74*), from 1.49% (without preservative) to 2.03% (glycerol) for Mg (*p = 0*.*99*) and from 0.53% (without preservative) to 0.85% (lyophilized cultures) for Ms (*p = 0*.*99*)

Significant differences were observed between mycoplasma strains with addition of complete medium (*p = 0*.*004*), horse serum (*p = 0*.*0001*), glycerol (*p = 0*.*00026*) and with lyophilised culture (*p = 0*.*00072*) (Table 1).

**Table 1.**
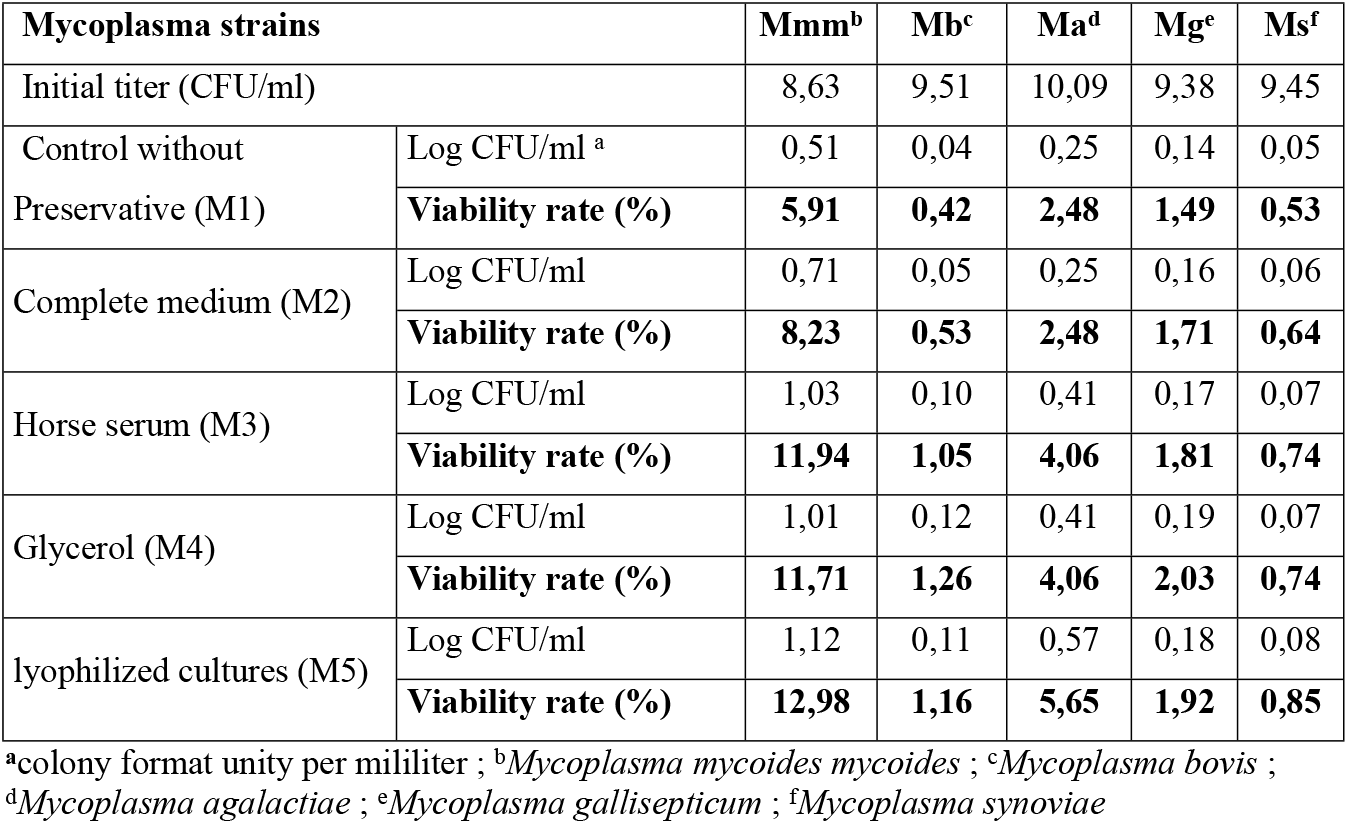
Viability of mycoplasma strains at + 37°C after 3 weeks

| Mycoplasma strains |  | Mmm<sup>b</sup> | Mb<sup>c</sup> | Ma<sup>d</sup> | Mg<sup>e</sup> | Ms<sup>f</sup> |
| --- | --- | --- | --- | --- | --- | --- |
| Initial titer (CFU/ml) |  | 8,63 | 9,51 | 10,09 | 9,38 | 9,45 |
| Control without Preservative (M1) | Log CFU/ml <sup>a</sup> | 0,51 | 0,04 | 0,25 | 0,14 | 0,05 |
|  | Viability rate (%) | **5,91** | **0,42** | **2,48** | **1,49** | **0,53** |
| Complete medium (M2) | Log CFU/ml | 0,71 | 0,05 | 0,25 | 0,16 | 0,06 |
|  | Viability rate (%) | **8,23** | **0,53** | **2,48** | **1,71** | **0,64** |
| Horse serum (M3) | Log CFU/ml | 1,03 | 0,10 | 0,41 | 0,17 | 0,07 |
|  | Viability rate (%) | **11,94** | **1,05** | **4,06** | **1,81** | **0,74** |
| Glycerol (M4) | Log CFU/ml | 1,01 | 0,12 | 0,41 | 0,19 | 0,07 |
|  | Viability rate (%) | **11,71** | **1,26** | **4,06** | **2,03** | **0,74** |
| lyophilized cultures (M5) | Log CFU/ml | 1,12 | 0,11 | 0,57 | 0,18 | 0,08 |
|  | Viability rate (%) | **12,98** | **1,16** | **5,65** | **1,92** | **0,85** |

<sup>a</sup>colony format unity per mililiter ; <sup>b</sup>*Mycoplasma mycoides mycoides* ; <sup>c</sup>*Mycoplasma bovis* ; <sup>d</sup>*Mycoplasma agalactiae* ; <sup>e</sup>*Mycoplasma gallisepticum* ; <sup>f</sup>*Mycoplasma synoviae*

### Viability of mycoplasma strains at +4°C

The refrigeration temperature of +4°C allowed for viable preservation of at least 3 months beyond which only the lyophilized cultures retained their viability with a loss of less than one (1) log CFU/ml (Fig 2).

**Fig 2.**
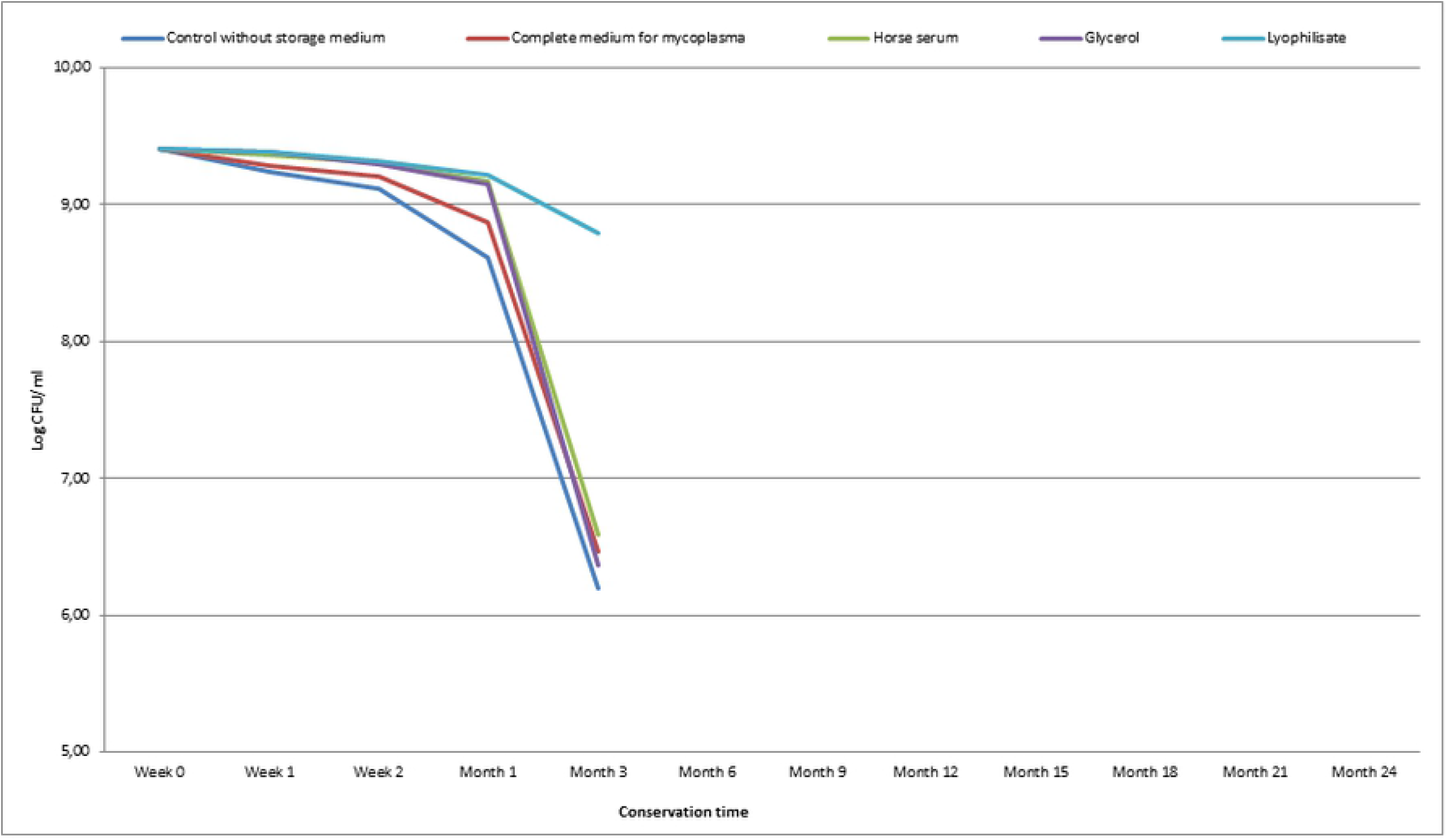
Mycoplasma viability at +4°C. Mean loss (in log) of the titre of 5 strains of mycoplasmas (Mmm, Mb, Ma, Mg, Ms) during 24 months of storage in different preservatives.

At +4°C, a significant effect (*p = 0*.*000417*) was observed with the storage media where the highest viability rate was obtained with the lyophilized cultures (17.81%) and the lowest with the glycerol (2.13%) after 6 months of storage.

The viability rates at 3 months of storage varied according to the mycoplasma strains and the storage media. A significant difference was observed with Mmm strain (0.46% and 17.85%, *p <0*.*001*), Mbovis strain (0.06% to 17.03%, *p <0*.*001*), the Ma strain (1.19 % to 28.24%, *p <0*.*001*), Mg strain (0.04% to 14.61%, *p <0*.*001*) and Ms strain (0.05% to 10.59%, *p < 0*.*001*)

The highest viability rates were obtained with the lyophilized cultures of Ma (28.24%), Mmm (17.85%), Mb (17.03%), Mg (14.61%) and Ms (10.59%); however, a viability of 5.91% was observed with the Mmm strain conserved in glycerol and 4.75% with the addition of the complete growth medium to the strain (Table 2).

**Table 2.** Viability of mycoplasma strains at + 4°C after 3 months

| **Mycoplasma strains** |  | **Mmm<sup>b</sup>** | **Mb<sup>c</sup>** | **Ma<sup>d</sup>** | **Mg<sup>e</sup>** | **Ms<sup>f</sup>** |
| --- | --- | --- | --- | --- | --- | --- |
| Initial titer (CFU/ml) |  | 8,63 | 9,51 | 10,09 | 9,38 | 9,45 |
| Control without preservative | Log CFU/ml <sup>a</sup> | 0,20 | 0,01 | 0,12 | 0,00 | 0,00 |
|  | **Viability rate (%)** | **2,32** | **0,06** | **1,19** | **0,04** | **0,05** |
| Complete medium | Log CFU/ml | 0,41 | 0,01 | 0,33 | 0,00 | 0,01 |
|  | **Viability rate (%)** | **4,75** | **0,09** | **3,27** | **0,04** | **0,07** |
| Horse serum | Log CFU/ml | 0,04 | 0,01 | 0,33 | 0,00 | 0,01 |
|  | **Viability rate (%)** | **0,46** | **0,11** | **3,27** | **0,01** | **0,1** |
| Glycerol | Log CFU/ml | 0,51 | 0,01 | 0,37 | 0,08 | 0,03 |
|  | **Viability rate (%)** | **5,91** | **0,11** | **3,67** | **0,85** | **0,32** |
| lyophilized cultures | Log CFU/ml | 1,54 | 1,62 | 2,85 | 1,37 | 1,00 |

|  | Viability rate (%) | 17,85 | 17,03 | 28,24 | 14,61 | 10,59 |
| --- | --- | --- | --- | --- | --- | --- |

<sup>a</sup>colony format unity per mililiter ; <sup>b</sup>*Mycoplasma mycoides mycoides* ; <sup>c</sup>*Mycoplasma bovis* ; <sup>d</sup>*Mycoplasma agalactiae* ; <sup>e</sup>*Mycoplasma gallisepticum* ; <sup>f</sup>*Mycoplasma synoviae*

### Viability of mycoplasma strains at −20°C

Overall, the mycoplasmas remained viable up to 24 months of storage at −20°C with an approximate titer drop of 0.01 log per titration session. The lyophilized cultures lost approximately 1 log at M24 post storage while with preservatives such as horse serum and glycerol the loss was 1 to 2 log CFU / ml (Fig 3).

**Fig 3.**
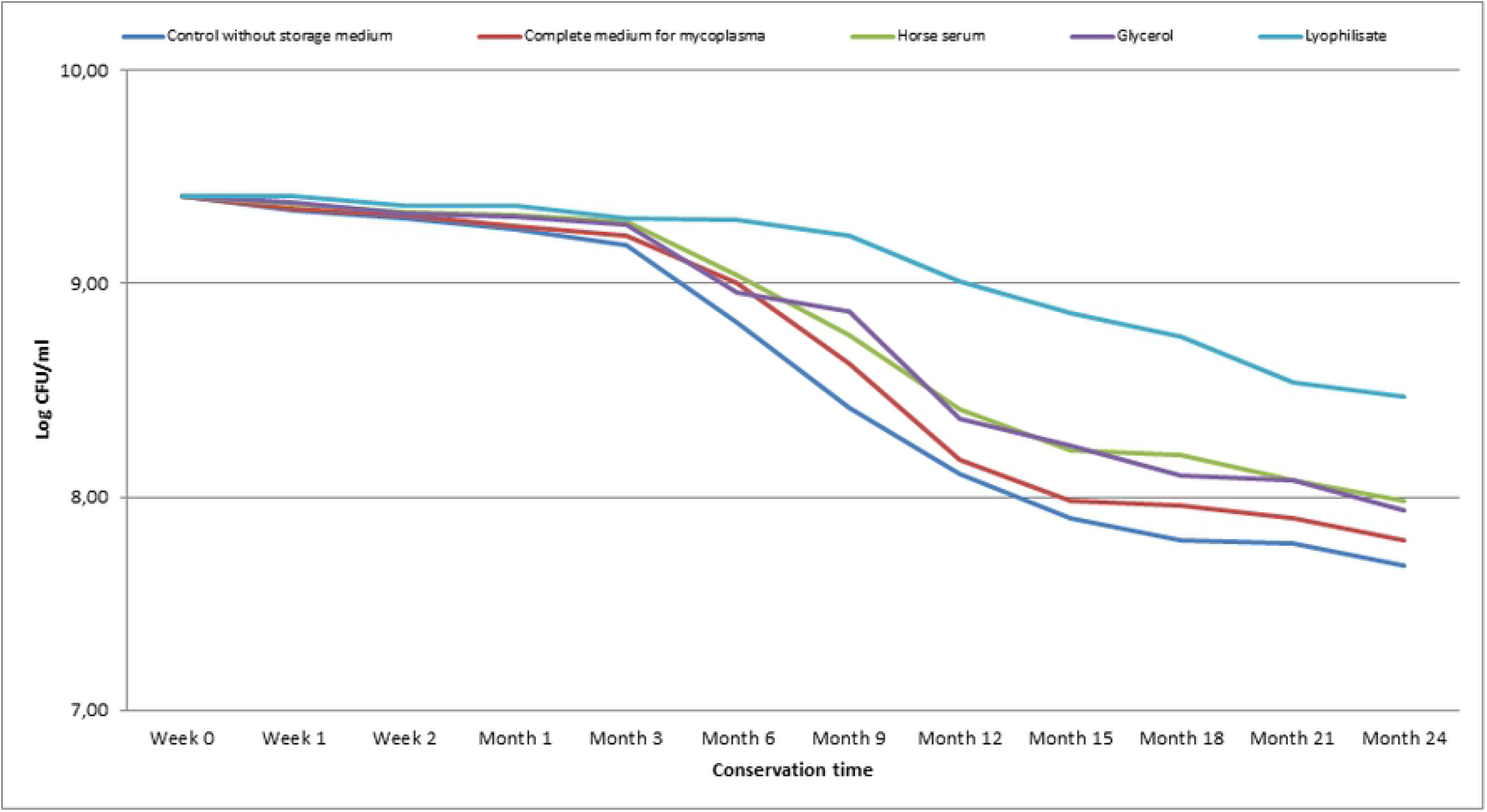
Mycoplasma viability at −20°C. Mean loss (in log) of the titre of 5 strains of mycoplasmas (Mmm, Mb, Ma, Mg, Ms) during 24 months of storage in different preservatives.

At −20°C, after 24 months of storage, viability rates varied by storage medium with 81.60% for the control without preservative, 82.88% with the complete medium, 84.79% with the addition of horse serum, at 84.36% with glycerol as preservatives and 90.04% with the lyophilized cultures. A non-significant effect (*p = 0*.*5229*) was observed between the viability rates of the mycoplasma strains and the storage media.

Viability rates varied by strain of mycoplasma and by preservative where lyophilized cultures, glycerol and horse serum maintained a rate above 90% viability. The viability was 92.73% to 99.34% with Mmm in horse serum and lyophilized cultures, 84.12% to 97.79% for Mb with complet medium and lyophilized cultures, 79.26% to 84.12% for Ma without preservative and lyophilized cultures, 76.76% to 83.15 % for Mg without preservative and lyophilized cultures and, 80.45% to 86.80% for Ms horse serum and lyophilized cultures (Table 3).

**Table 3.** Viability of mycoplasma strains at −20°C after 24 months

| Mycoplasma strains |  | Mmm<sup>b</sup> | Mb<sup>c</sup> | Ma<sup>d</sup> | Mg<sup>e</sup> | Ms<sup>f</sup> |
| --- | --- | --- | --- | --- | --- | --- |
| Initial titer |  | 8,63 | 9,51 | 10,09 | 9,38 | 9,45 |
| Control without preservative | Log CFU/ml <sup>a</sup> | 7,20 | 8,10 | 8,00 | 7,20 | 7,90 |
|  | Viability rate (%) | 83,46 | 85,17 | 79,26 | 76,76 | 83,62 |
| Complete medium | Log CFU/ml | 7,40 | 8,00 | 8,10 | 7,80 | 7,70 |
|  | Viability rate (%) | 85,77 | 84,12 | 80,25 | 83,15 | 81,51 |
| Horse serum | Log CFU/ml | 8,00 | 8,20 | 8,40 | 7,70 | 7,60 |
|  | Viability rate (%) | 92,73 | 86,22 | 83,22 | 82,09 | 80,45 |
| Glycerol | Log CFU/ml | 8,10 | 8,20 | 8,30 | 7,50 | 7,60 |
|  | Viability rate (%) | 93,89 | 86,22 | 82,23 | 79,96 | 80,45 |
| lyophilized cultures | Log CFU/ml | 8,57 | 9,30 | 8,49 | 7,80 | 8,20 |
|  | Viability rate (%) | 99,34 | 97,79 | 84,12 | 83,15 | 86,80 |

<sup>a</sup>colony format unity per mililiter ; <sup>b</sup>*Mycoplasma mycoides mycoides* ; <sup>c</sup>*Mycoplasma bovis* ;
<sup>d</sup>*Mycoplasma agalactiae* ; <sup>e</sup>*Mycoplasma gallisepticum* ; <sup>f</sup>*Mycoplasma synoviae*

### Viability of mycoplasma strains at −85 ° C

After 24 months of storage at −85°C, the mean losses of titer of mycoplasmas in the whole were from 0.8 log (lyophilized cultures) to 1.06 log (horse serum, glycerol), the complete medium and controls with mean losses of 1.23 and 1.31 log CFU / ml. Cryopreservatives (glycerol and horse serum) and lyophilized cultures made it possible to maintain the mycoplasma strains viable with progressive losses over time (Fig 4).

**Fig 4.**
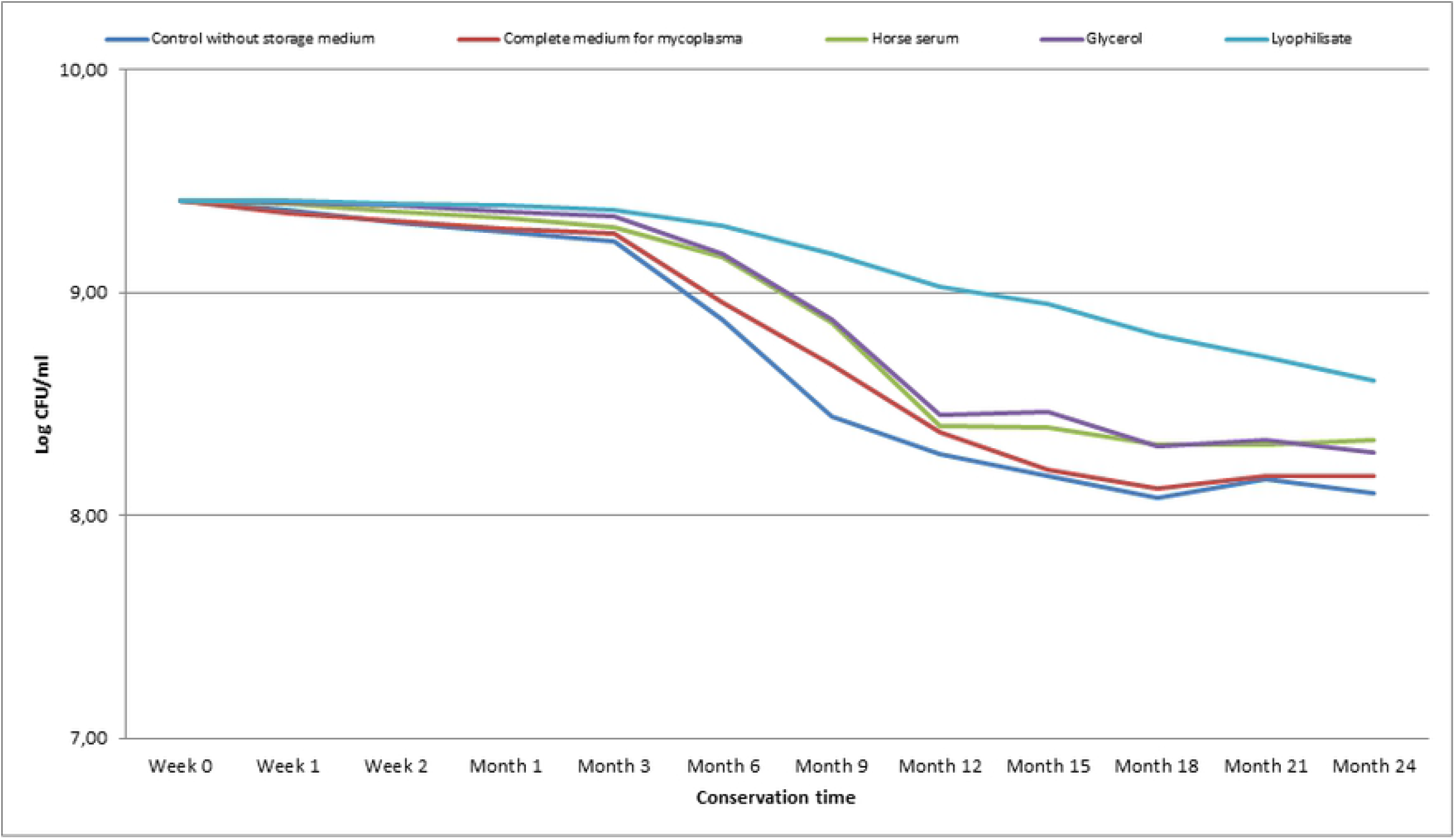
Mycoplasma viability at −85°C. Mean loss (in log) of the titre of 5 strains of mycoplasmas (Mmm, Mb, Ma, Mg, Ms) during 24 months of storage in different preservatives.

At −85°C, non-significant overall viability rates (*p-value = 0*.*8035*) were obtained between the mycoplasma strains and the preservatives at 24 months of storage with respectively overall average viability rates of 86.06%, 86.91%, 88.61%, 87.98% and 91.44% with control without preservative, complete medium, horse serum, glycerol and lyophilized cultures.

Viability rates of the mycoplasma strains after 24 months of storage under −85°C depending on the preservatives were from 92.73% (without preservatives) to 97.71% (lyophilized cultures) for the Mmm strain, 86.22 to 92.53% for Mb glycerol and lyophilized cultures, from 80.25 to 92.14% for Ma without preservative and lyophilized cultures, from 85.29 to 88.48% for Mg and from 85.74% to 86.8% for Ms (Table 4).

**Table 4.**
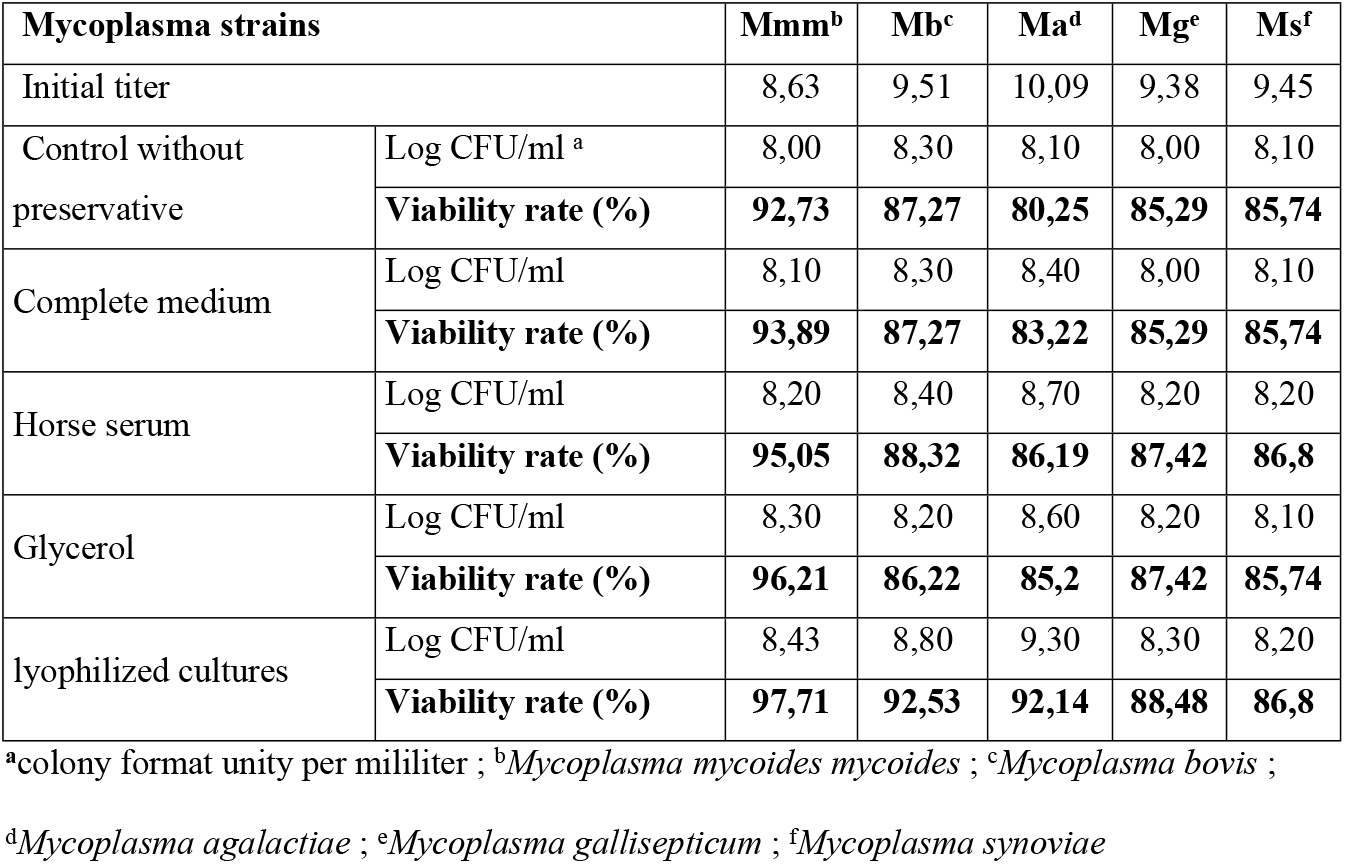
Viability of mycoplasma strains at −85°C after 24 months

| **Mycoplasma strains** |  | **Mmm<sup>b</sup>** | **Mb<sup>c</sup>** | **Ma<sup>d</sup>** | **Mg<sup>e</sup>** | **Ms<sup>f</sup>** |
| --- | --- | --- | --- | --- | --- | --- |
| Initial titer |  | 8,63 | 9,51 | 10,09 | 9,38 | 9,45 |
| Control without preservative | Log CFU/ml <sup>a</sup> | 8,00 | 8,30 | 8,10 | 8,00 | 8,10 |
|  | **Viability rate (%)** | **92,73** | **87,27** | **80,25** | **85,29** | **85,74** |
| Complete medium | Log CFU/ml | 8,10 | 8,30 | 8,40 | 8,00 | 8,10 |
|  | **Viability rate (%)** | **93,89** | **87,27** | **83,22** | **85,29** | **85,74** |
| Horse serum | Log CFU/ml | 8,20 | 8,40 | 8,70 | 8,20 | 8,20 |
|  | **Viability rate (%)** | **95,05** | **88,32** | **86,19** | **87,42** | **86,8** |
| Glycerol | Log CFU/ml | 8,30 | 8,20 | 8,60 | 8,20 | 8,10 |
|  | **Viability rate (%)** | **96,21** | **86,22** | **85,2** | **87,42** | **85,74** |
| lyophilized cultures | Log CFU/ml | 8,43 | 8,80 | 9,30 | 8,30 | 8,20 |
|  | **Viability rate (%)** | **97,71** | **92,53** | **92,14** | **88,48** | **86,8** |

<sup>a</sup>colony format unity per mililiter ; <sup>b</sup>*Mycoplasma mycoides mycoides* ; <sup>c</sup>*Mycoplasma bovis* ;
<sup>d</sup>*Mycoplasma agalactiae* ; <sup>e</sup>*Mycoplasma gallisepticum* ; <sup>f</sup>*Mycoplasma synoviae*

Significant differences in viability rates of Mmm, Mb, Ma, Mg and Ms were observed between different preservatives and temperature conditions. In general, all additives improved the viability rate of mycoplasmas compared to the control group. The highest viability rates of Mmm, Ma, Mg and Ms were observed at −85°C with lyophilized cultures while for Mb this rate was observed with glycerol at −85°C.

## Discussion

The viability of vaccine and wild strains of mycoplasmas depends on the storage conditions, in particular temperature, preservatives and shelf life. With regards to temperature, to maintain cell cultures viability for extended periods, storage at −70 to −85°C was much more preferable especially with the addition of cryopreservatives such as dimethyl sulfoxide (DMSO) or glycerol. Our results showed that storage at −85°C was preferable than at −20°C which despite the loss of viability of the mycoplasmas during the freezing and thawing process seems better than storage at +4°C intended for short-term storage (not more than 3 months). The lyophilized cultures of mycoplasmas allowed a better preservation of the viability whatever the temperature (+4, −20, −85°C), which was also observed that the viability of the lyophilized cultures of mycoplasmas was stable for at least 34 months at + 4°C. In general, lyophilized cultures of various mycoplasmas were well preserved after long-term storage at 4°C [15]. It is a fact that, at −85°C, the metabolism process is completely stopped unlike −20°C where the metabolism continues but at a very slow rate. Warming during thawing of strains affected negatively on the viability rate of frozen mycoplasmas. Rapid warming of frozen strains is preferable because it apparently prevents recrystallization of intracellularice. Cryoprotective agents such as glycerol have been shown to be the most effective in protecting mycoplasmas cultures against freezing loss during the cooling process, possibly by reducing the fraction of frozen cellular water [16]. The effect of freeze-thaw conditions on the viability of Mb is a significant challenge for the preservation of these bacteria. The lack of peptidoglycan cell wall in mycoplasmas makes them susceptible to ice crystal formation during freeze / thaw processes [17]. It emerged from this study that glycerol led to an improvement in the survival of Mycoplasmas due to its bacteriostatic activity contributing to the inhibition of other bacterial proliferation [18]. The Mycoplasma strains tested in this study showed varying degrees of sensitivity to temperatures such as +37°C, +4°C, −20°C and −85°C. Cultures die quickly at +37°C and at room temperature which would be incubation and not storage temperatures and at +4°C where they would survive for 2 weeks in solid medium but less in liquid; on the other hand at −20°C or less, the cultures will be able to survive for 6 to 12 months [19]. The viability of the microorganisms would be important with liquid nitrogen [20] but in the framework of routine diagnosis of mycoplasmosis from samples, the critical factors would be the storage temperature, the duration and the type of swab for the isolation of strains of *Mycoplasma gallisepticum* and *Mycoplasma synoviae* [21]. Compared to *Mycoplasma bovis*, its viability in milk would depend on the storage temperature which for 5 days at +4°C would reduce the number but the addition of glycerol (10 - 30%) to the frozen milk samples improved the survival of *Mycoplasma bovis* whose detection will be maximized in fresh milk grown without storage [22]. The duration of viability of the mycoplasma strains under refrigeration temperature (+4°C) did not exceed 3 months, unlike at temperatures of −20°C and −85°C; this was confirmed by the study performed under anaerobic conditions where viable mycoplasmas were found with little or no reduction in titers after storage for 8 weeks at −30°C with a reduction in titers of 3 log10 produced after 4 weeks of storage at +4°C [23]. Freezing for 14 weeks did not significantly reduce *Mycoplasma bovis* titer compared to freezing for 2 weeks, and a second freeze-thaw cycle reduced *Mycoplasma bovis* count by approximately 0.5 log compared to a single freeze-thaw cycle [24]. Lyophilized cultures of Mycoplasma spp. stored at +4° C had remained viable for 18 to 22 years and 82% of Mycoplasmas initially lyophilized then reconstituted were viable after 16 years of storage at −70°C [25]. Glycerol, in addition to its bacteriostatic power, is the main source of carbon and energy, especially for bacteria without cell walls of the genus Mycoplasma [26].

## Conclusion

This study revealed that the lyophilized mycoplasmas without stabilizer, withstood storage at +4°C for more than 3 months and that mycoplasma broths supplemented in storage medium (complete mycoplasma medium, horse sera, glycerol) could not remain viable beyond 3 months. It should also be noted that the addition of cryopreservatives such as glycerol to the cultures improved the viability of mycoplasmas. Temperatures of +37°C and +4°C were respectively for incubation (3 weeks) and short-term storage temperatures (3 months); on the other hand, for long-term storage the temperature of −85°C is much more advantageous than − 20°C (>24 months in this study).

## Acknowledgements

The study was carried out with the financial support of the Malian government through “competitive funds for research and technological innovation” and the technical support of scientific partners such as the International Atomic Energy Agency (IAEA) Vienna-Austria and the Center for International Cooperation in Agricultural Research for Development (CIRAD) Montpellier-France.

## Author Contributions

Conceptualization: SERY A., SIDIBE C.A.K., NIANG M.

Formal analysis: SERY A., SIDIBE C.A.K., NIANG M.

Funding acquisition: SERY A.

Investigation: SERY A., SIDIBE C.A.K., KONE M., SACKO B., NIANG M.

Methodology: SERY A., SIDIBE C.A.K., KONE M., SACKO B., NIANG M.

Supervision: SERY A., SIDIBE C.A.K., NIANG M.

Validation: SERY A., SIDIBE C.A.K., NIANG M.

Visualization: SERY A., SIDIBE C.A.K., NIANG M.

Writing – original draft: SERA A.

Writing – review & editing: SERY A., SIDIBE C.A.K., NIANG M, AWUNI J, AMANFU WILLIAM

## References

1. Aebi M, Bodmer M, Frey J, Pilo P (2012) Herd-specific strains of Mycoplasma bovis in outbreaks of mycoplasmal mastitis and pneumonia. Vet Microbiol 157:363–368. https://doi.org/10.1016/j.vetmic.2012.01.006

2. Aebi M, van den Borne BHP, Raemy A, et al (2015) Mycoplasma bovis infections in Swiss dairy cattle: a clinical investigation. Acta Vet Scand 57:10. https://doi.org/10.1186/s13028-015-0099-x

3. Arcangioli MA, Chazel M, Sellal E, et al (2011) Prevalence of Mycoplasma bovis udder infection in dairy cattle: preliminary field investigation in southeast France. N Z Vet J 59:75–78. https://doi.org/10.1080/00480169.2011.552856

4. Castillo-Alcala F, Bateman KG, Cai HY, et al (2012) Prevalence and genotype of Mycoplasma bovis in beef cattle after arrival at a feedlot. Am J Vet Res 73:1932–1943. https://doi.org/10.2460/ajvr.73.12.1932

5. Gautier-Bouchardon AV (2018) Antimicrobial Resistance in Mycoplasma spp. Microbiol Spectr 6:. https://doi.org/10.1128/microbiolspec.ARBA-0030-2018

6. Hazelton MS, Morton JM, Bosward KL, et al (2018) Isolation of Mycoplasma spp. and serological responses in bulls prior to and following their introduction into Mycoplasma bovis-infected dairy herds. J Dairy Sci 101:7412–7424. https://doi.org/10.3168/jds.2018-14457

7. Addey J, Taylor-Robinson D, Dimic M (1970) Viability Of Mycoplasmas After Storage In Frozen Or Lyophilised States. J Med Microbiol 3:137–45. https://doi.org/10.1099/00222615-3-1-137

8. Amores J, Sánchez A, Martín AG, et al (2010) Viability of Mycoplasma agalactiae and Mycoplasma mycoides subsp. capri in goat milk samples stored under different conditions. Vet Microbiol 145:347–350. https://doi.org/10.1016/j.vetmic.2010.03.030

9. Al-Farha AA-B, Khazandi M, Hemmatzadeh F, et al (2018) Evaluation of three cryoprotectants used with bovine milk affected with Mycoplasma bovis in different freezing conditions. BMC Res Notes 11:216. https://doi.org/10.1186/s13104-018-3325-6

10. Biddle MK, Fox LK, Hancock DD, et al (2004) Effects of Storage Time and Thawing Methods on the Recovery of Mycoplasma Species in Milk Samples from Cows with Intramammary Infections. J Dairy Sci 87:933–936. https://doi.org/10.3168/jds.S0022-0302(04)73237-3

11. Christensen N, Yavari C, McBain A, Bradbury J (1994) Investigations into the survival of Mycoplasma gallisepticum, Mycoplasma synoviae and Mycoplasma iowae on materials found in the poultry house environment. Avian Pathol J WVPA 23:127–43. https://doi.org/10.1080/03079459408418980

12. Gourlay RN, Howard CJ, Thomas LH, Wyld SG (1979) Pathogenicity of some Mycoplasma and Acholeplasma species in the lungs of gnotobiotic calves. Res Vet Sci 27:233–237

13. Hayflick L, Stanbridge E (1967) Isolation and Identification of Mycoplasma from Human Clinical Materials*. Ann N Y Acad Sci 143:608–621. https://doi.org/10.1111/j.1749-6632.1967.tb27705.x

14. Frey ML, Hanson RP, Andrson DP (1968) A medium for the isolation of avian mycoplasmas. Am J Vet Res 29:2163–2171

15. Raccach M, Rottem S, Razin S (1975) Survival of frozen mycoplasmas. Appl Microbiol 30:167–171

16. Miller RH, Mazur P (1976) Survival of frozen-thawed human red cells as a function of cooling and warming velocities. Cryobiology 13:404–414. https://doi.org/10.1016/0011-2240(76)90096-1

17. Pegg DE (2007) Principles of cryopreservation. Methods Mol Biol Clifton NJ 368:39–57. https://doi.org/10.1007/978-1-59745-362-2_3

18. Roger V, Fonty G, Andre C, Gouet P (1992) Effects of glycerol on the growth, adhesion, and cellulolytic activity of rumen cellulolytic bacteria and anaerobic fungi. Curr Microbiol 25:197–201. https://doi.org/10.1007/BF01570719

19. Klieneberger-Nobel E (1962) Pleuropneumonia-like Organisms (PPLO) Mycoplasmataceae. Academic Press

20. Bajerski F, Bürger A, Glasmacher B, et al (2020) Factors determining microbial colonization of liquid nitrogen storage tanks used for archiving biological samples. Appl Microbiol Biotechnol 104:131–144. https://doi.org/10.1007/s00253-019-10242-1

21. Ball C, Felice V, Ding Y, et al (2020) Influences of swab types and storage temperatures on isolation and molecular detection of Mycoplasma gallisepticum and Mycoplasma synoviae. Avian Pathol J WVPA 49:106–110. https://doi.org/10.1080/03079457.2019.1675865

22. Boonyayatra S, Fox LK, Besser TE, et al (2010) Effects of storage methods on the recovery of Mycoplasma species from milk samples. Vet Microbiol 144:210–213. https://doi.org/10.1016/j.vetmic.2009.12.014

23. Cheng H-S, Shen C-W, Wang S-R (2007) Effect of storage conditions on detection of mycoplasma in biopharmaceutical products. In Vitro Cell Dev Biol Anim 43:113–9. https://doi.org/10.1007/s11626-007-9015-7

24. Gille L, Boyen F, Van Driessche L, et al (2018) Short communication: Effect of freezer storage time and thawing method on the recovery of Mycoplasma bovis from bovine colostrum. J Dairy Sci 101:609–613. https://doi.org/10.3168/jds.2017-13130

25. Furr PM, Taylor-Robinson D (1990) Long-term viability of stored mycoplasmas and ureaplasmas. J Med Microbiol 31:203–206. https://doi.org/10.1099/00222615-31-3-203

26. Blötz C, Stülke J (2017) Glycerol metabolism and its implication in virulence in Mycoplasma. FEMS Microbiol Rev 41:640–652. https://doi.org/10.1093/femsre/fux033

